# Putative G-quadruplex forming sequence signatures in genes differentially transcribed upon loss of BLM or WRN helicases

**DOI:** 10.1101/013664

**Authors:** John Smestad, L. James Maher

**Affiliations:** Mayo Clinic Medical Scientist Training Program, Mayo Clinic College of Medicine, 200 First St. SW, Rochester, MN 55905, USA; Department of Biochemistry and Molecular Biology, Mayo Clinic College of Medicine, 200 First St. SW, Rochester, MN 55905, USA

## Abstract

Putative G-quadruplex-forming sequences (PQS) have long been implicated in regulation of DNA replication and transcription, though their actual roles are unknown. To gain insight into potential PQS transcriptional function, we map and analyze PQS motifs in promoters of genes differentially-expressed in Bloom Syndrome (BS) and Werner Syndrome (WS), two human genetic disorders resulting in loss of PQS-interacting RecQ helicases. Non-B-DNA structures at PQS might be stabilized in these syndromes. For BS and WS we demonstrate that PQS promoter abundance is generally higher in down-regulated genes and lower in up-regulated genes, and show that these effects are position-dependent. To interpret these correlations we determined genome-wide PQS correlations with transcription using epigenetic information to predict gene expression. We report that 33% and 35% of analyzed PQS positions in promoter antisense and sense strands, respectively, displayed statistically-significant correlation with gene expression. Of these statistically-significant positions, 100% and 84% on antisense and sense strands, respectively, were correlated with reduced expression. This suggests that promoter PQS repress transcription. Finally, we report neural network clustering analysis of PQS motifs to demonstrate that genes differentially-expressed in BS and WS are significantly biased in their PQS motifs, suggesting an unappreciated biological relationship between PQS, RecQ helicases, and transcription.

**REVIEWER LINKS TO DEPOSITED DATA:** ftp://www.jsmes.net/PQS_Genomics

## INTRODUCTION

Bloom Syndrome (BS) and Werner Syndrome (WS) are human disorders resulting from loss-of-function mutations in DNA helicases belonging to the RecQ helicase family (Bloom 1954; Epstein et al. 1966; German et al. 1979; German 1993; Yu et al. 1996; Lauper et al. 2013). BS patients present with short stature, immunodeficiency, and photo-sensitivity. WS is classically associated with progeria. Both syndromes result in decreased fertility and increased carcinogenesis. The BLM and WRN helicases implicated in BS and WS, respectively, have both been shown to unfold G-quadruplex structures assembled *in vitro* (Sun et al. 1998; Fry and Loeb 1999). It has been proposed that these helicases function *in vivo* by resolving intrastrand and interstrand G-quadruplexes hypothesized to form at putative G-quadruplex-forming sequences (PQS) during homologous recombination (Harmon et al. 1999), base-excision repair (WRN only) (Ahn et al. 2004), in telomeres during cellular replication (Crabbe et al. 2004; Opresko et al. 2005; Crabbe et al. 2007), and in regulation of gene transcription (Johnson et al. 2010). Here PQS are identified by an algorithm (see Methods) analyzing the potential to form intrastrand G-quadruplexes under proper conditions if DNA strands were separated (Huppert and Balasubramanian 2005).

Analysis of transcriptional perturbations in BS and WS has identified effects on various cellular signaling pathways including control of growth/proliferation, death/survival, protein synthesis, gene expression, and development (Cheung et al. 2014; Nguyen et al. 2014). Interestingly, it has also been noted that genes differentially expressed in BS and WS have increased PQS abundance, suggesting that transcriptional changes upon loss of RecQ helicases could result from failure to properly suppress G-quadruplex structures (Johnson et al. 2010; Nguyen et al. 2014). Despite the absence of conclusive biochemical evidence for G-quadruplex structures at PQS in these genes, sequence-expression correlations are compelling.

Little is yet known concerning specific PQS motifs in the promoters of genes differentially sensitive to loss of BLM and WRN helicases. Such knowledge might provide insight into how PQS /RecQ helicase interactions modulate gene expression. In the current work, we therefore analyze the abundance and sequences of PQS within 2 kbp of transcription start sites (TSS) for genes differentially expressed in BS and WS. We elucidate statistically significant PQS abundance patterns in these genes vs. genes whose transcription is not altered by RecQ helicase loss. We also use a new approach to correlate PQS location with transcriptional activation or repression. The method applies epigenetic information to predict gene expression, with subsequent analysis of the modeling error and correlation with PQS position. These two methods map intrinsic PQS transcription regulatory effects, and predict how PQS abundance at discrete positions correlates with transcriptional changes upon BLM or WRN helicase loss. Finally, we analyze PQS motifs using neural network clustering to demonstrate that genes differentially-expressed in BS and WS are significantly biased in their PQS motifs. This suggests an unappreciated biological relationship between PQS, RecQ helicases, and transcription.

## RESULTS

### PQS abundance in genes differentially expressed in BS and WS

We hypothesized that PQS abundance in promoters of genes differentially expressed in BS and WS reflects transcriptional sensitivity to RecQ helicase activity. We used published gene expression array datasets comparing BS and WS patient fibroblasts to normal control fibroblasts to identify genes that are significantly up-and down-regulated in BS and WS [absolute fold expression change ≥ 1.5 with an adjusted p-value < 0.05 and false discovery rate (FDR) < 0.1 for BS genes and FDR < 0.05 for WS genes] (Cheung et al. 2014; Nguyen et al. 2014). The BS dataset consisted of 1010 up-regulated genes and 141 down-regulated genes (Nguyen et al. 2014), and the WS dataset consisted of 1046 up-regulated genes and 540 down-regulated genes. Comparing genes identified as differentially-expressed in each syndrome, we find them to be largely non-overlapping (**Supplemental Figure S1**). Notably, the fraction of these differentially-expressed genes that contains at least one PQS within 2 kbp of a TSS (BS: 84% of up-regulated genes and 90% of down-regulated genes, WS: 74% of up-regulated genes and 84% of down-regulated genes), is high compared to the genomic average of 55% (**Supplemental Table S1**).

We analyzed how PQS abundance varies as a function of position and strand (sense or antisense) near promoters of genes differentially expressed in BS and WS, compared to other genes. Histograms showing PQS abundance (raw counts) as a function of position upstream and downstream of the TSS on sense and antisense strands are shown in **Supplemental Figure S2**. We then compared PQS abundance between genes differentially expressed in BS and WS vs. all other genes. A heat map showing PQS abundance (raw counts) as a function of position, normalized to total number of TSS in the respective datasets, is presented in **Figure 1A**. We also conducted quantitative analysis of PQS abundance in genes differentially expressed in BS and WS vs. all other genes, using 200 bp bins, repeated at a 10-bp interval, spanning 2 kbp upstream and downstream of known TSS on both strands. For each bin, 1-way ANOVA p-values were calculated comparing PQS abundance in the differentially-expressed gene dataset to all other genes. The statistically-significant result, shown in **Figure 1B**, is intriguing. Genes up-and down-regulated in BS and WS tend to have opposite patterns of PQS abundance near TSS. Relative to the remainder of the genome, genes up-regulated in BS and WS tend to have fewer PQS, while genes down-regulated in BS and WS tend to have more PQS. An exception to this pattern is that BS up-regulated genes have more PQS between 160 and 680 bp downstream of the TSS on the sense strand. Other PQS abundance patterns are listed in **Supplemental Table S2**. Threshold p-values and FDR for the analysis, determined by statistical simulation with randomly-generated gene sets, are presented in **Supplemental Table S3**.

**Figure 1.**
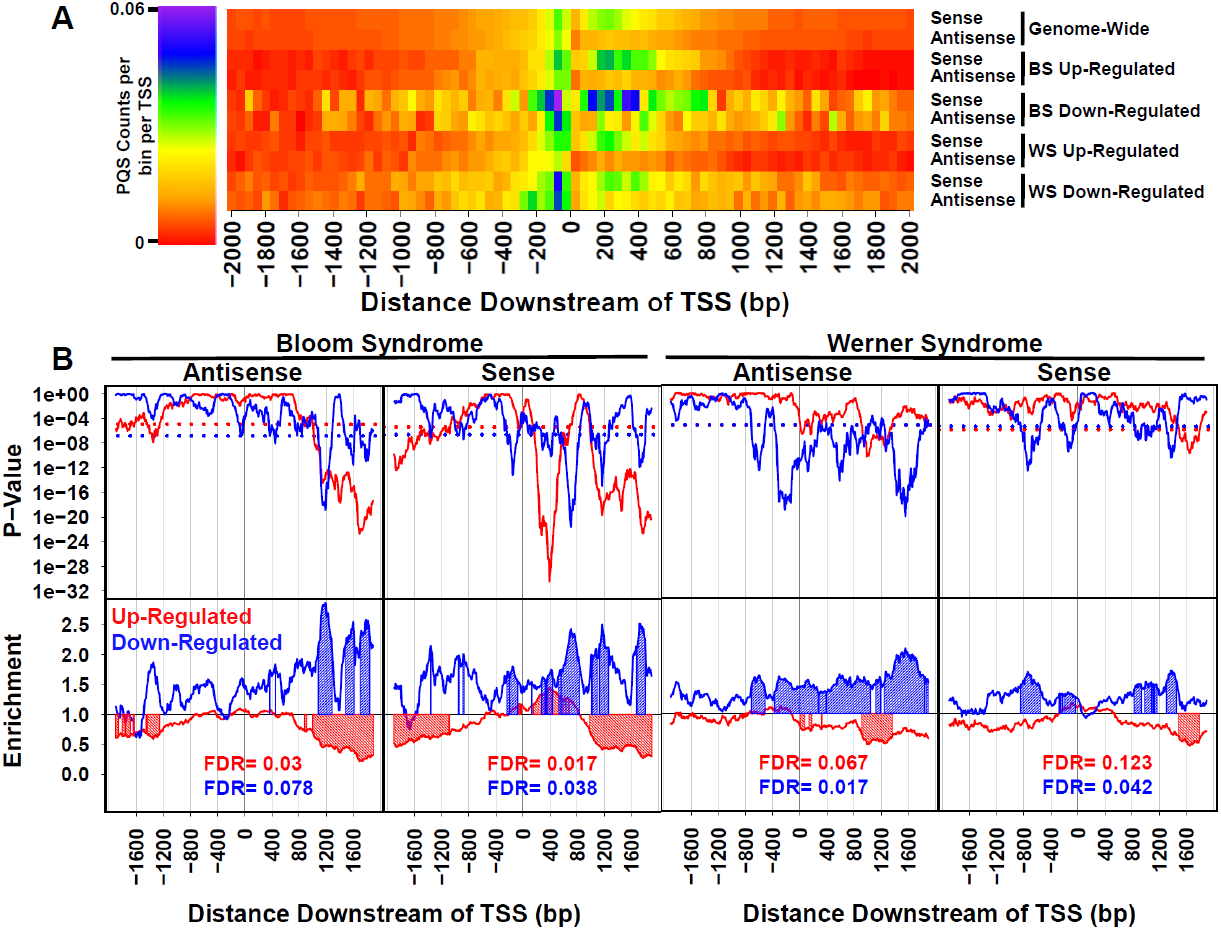
PQS occurrence in genes differentially expressed in BS and WS. PQS occurrence in sense (S) and antisense (AS) strands was analyzed independently. (A) Heat map showing PQS occurrence genome-wide and in genes differentially-expressed in BS and WS, represented as raw counts normalized to total number of TSS in each dataset. (B) Top panels show p-values for comparing PQS abundance per TSS between genes differentially expressed in BS and WS and all other genes. Dotted lines represent p-value cutoffs for determining statistical significance, with less than 1% of data points from a random gene dataset of the same size as the test dataset having a p-value below this threshold. Bottom panels show PQS enrichment ratio in genes differentially expressed in BS and WS, with values > 1 indicating that PQS are more abundant in the differentially-expressed gene set. Both p-values and enrichment ratios were calculated using a 200 bp bin value repeated at a 10 bp interval. Regions with shaded peaks represent locations of statistically-significant PQS excess or scarcity.

### Correlation of PQS patterns and transcription

To interpret PQS abundance for genes differentially expressed in BS and WS, we studied how PQS location relative to the TSS correlates with gene expression for all human genes. A previous study correlated PQS position and gene transcription while controlling for gene family, function, and promoter similarity (Du et al. 2008). In that work, PQS up to 500 bp downstream of TSS were correlated with increased gene expression. PQS up to 500 bp upstream of the TSS were not significantly correlated with gene expression. In our analysis, positions found to have altered PQS abundance in genes differentially expressed in BS and WS often occur more than 500 bp from TSS (86% for BS, 71% for WS). No data were available for genome-wide correlation between transcription and PQS in such positions.

We therefore implemented a novel analysis combining published epigenetic and gene expression data from 7 human cell lines (H1-hESC, HeLa-S3, K562, HUVEC, HepG2, NHEK, and GM12878) to correlate PQS positions within 2 kbp of TSS genome-wide with gene expression. The method successfully predicts gene expression using epigenetic data, and sorts calculated prediction errors (actual expression minus predicted expression) by PQS position near TSS. The method is unique in its robust statistics that control for gene epigenetic signature to isolate regulatory effects not detectable by standard association-based methods. Our approach to modeling gene expression was inspired by a prior publication (Dong et al. 2012). Bayesian linear regression models were trained on TSS-specific cap analysis of gene expression (CAGE) measurements, epigenetic data including abundance of local histone modifications (H3K4me1, H3K4me2, H3K4me3, H3K9ac, H3K27ac, H3K27me3, H3K36me3, H3K79me2, H4K20me1), histone variant H2A.Z, and GC fraction 5 kbp upstream of the TSS. For each histone modification and variant, we identified the position near the TSS where the epigenetic signature best correlated with gene expression (**Figure 2A**, **Supplemental Figure S3**). Model training was conducted using half of the PQS-free TSS (i.e.no PQS within 2 kbp of the TSS) for each cell line. Resulting models were then applied to the remaining TSS in the dataset (the remaining half of all PQS-free TSS as well as all PQS-containing TSS) to generate gene expression predictions (**Supplemental Tables S4, S5, and S6**). Thus, each TSS in the test dataset has both predicted and measured transcription values. A representative plot of predicted vs. measured transcription is shown in **Figure 2B**. The difference between measured and predicted transcription is the prediction error. Prediction error values > 0 indicate greater-than-predicted transcription, and values < 0 indicate less-than-predicted transcription. Prediction errors for genes with ≥ 1 PQS within 2 kbp of the TSS were sorted based on PQS position and this distribution compared to the distribution of prediction errors for PQS-free TSS using 1-way ANOVA. Statistical methods were used to determine threshold p-values and to estimate the FDR by repetition of the analysis after replacement of the prediction error values with randomly sampled values from the control distribution. A representative plot for this type of analysis is shown in **Figure 2C**.

**Figure 2.**
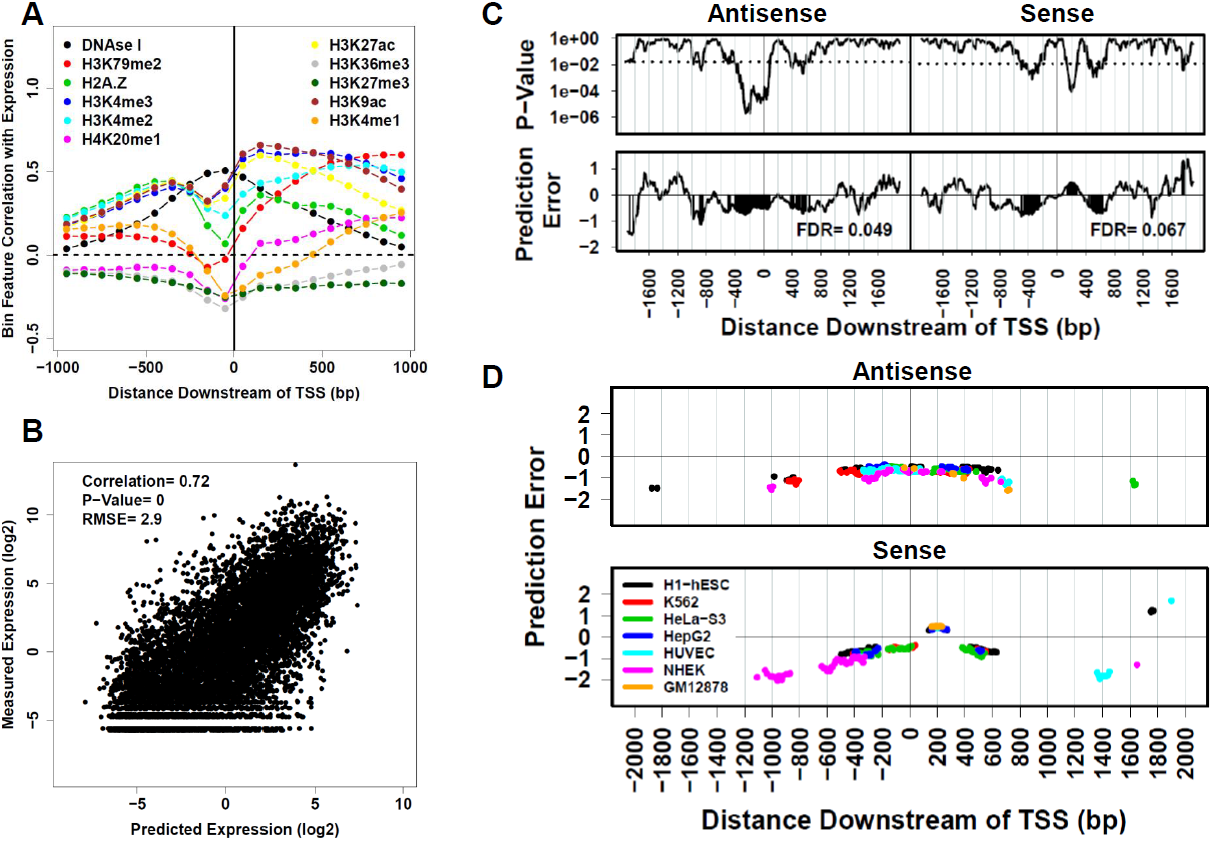
Epigenetic prediction of gene expression to identify PQS positions correlated with altered gene expression. (A) Correlation of log_2_-transformed epigenomic track signals with log_2_-transformed gene expression represented as a function of location 1 kbp upstream and downstream of TSS. Bin size is 100 bp. Prior to log_2_-transformation, 0.25% of the maximal bin value was added to each bin feature to avoid the log_2_(0) issue. (B) Representative correlation of CAGE TSS-specific gene expression measurements with epigenetic model predictions generated through Bayesian linear regression. Models were trained on datasets containing half of all PQS-free TSS in the human genome. Gene expression predictions were then generated for the remainder of the PQS-free TSS. (C) Analysis of model prediction error as a function of PQS position, assigned to sense (S) and antisense (AS) strand effects. Prediction error estimates were sorted into 200-bp bins iterated at a 10-bp interval, based on the position of the PQS nearest to the TSS, and then compared to the prediction error estimates for PQS-free TSS using 1-way ANOVA. The top panels show the p-value from this analysis as a function of position with respect to the TSS. The dashed line represents the p-value threshold used for determining statistical significance in the analysis. The bottom panels show prediction error averages for the binned values. Areas with shaded peaks represent locations where the p-value is statistically significant for rejection of the null hypothesis. Prediction error value > 0 represent positions where PQS presence correlates with higher gene expression. Prediction error values < 0 represent positions where PQS presence correlates with lower gene expression. (D) Aggregate map of all statistically-significant positions from prediction error analysis based on seven human cell lines.

This analysis was repeated for each of 7 human cell lines, generating a composite analysis of all statistically-significant PQS positions correlated with gene expression (**Figure 2D**). This is the first PQS correlation analysis incorporating epigenetic data from multiple cell lines and extending the study region to 2 kbp upstream and downstream of TSS. Interestingly, the general agreement for data from different cell lines suggests that PQS position correlates with gene expression regardless of epigenetic context. The analysis correlates PQS position in the antisense strand with lower gene expression, regardless of PQS position upstream or downstream of the TSS. In contrast, PQS in the sense strand show different correlations depending on position. PQS in the sense strand are correlated with lower gene expression, except for PQS positioned downstream of the TSS between 140-270 bp, 1750-1770 bp, and 1900 bp. PQS at these three sense strand positions were correlated with increased gene expression. This analysis did not find PQS correlations with gene expression at all locations. In total, 33% and 35% of analyzed PQS positions in the antisense and sense strands, respectively, displayed statistically-significant PQS abundance correlated with gene expression. These data represent a large improvement over previous studies in terms of resolution and coverage.

It was then possible to compare position-dependent PQS correlations with gene expression for all genes to data for genes differentially expressed in BS and WS (**Figure 3**). For both DNA strands of genes up-regulated in BS and WS, TSS-proximal PQS are underrepresented at positions correlated with low gene expression. Similarly, for both DNA strands of genes down-regulated in BS and WS, TSS-proximal PQS are over-represented at positions correlated with low gene expression. These observations are together consistent with a model in which the BLM and WRN helicases suppress some inhibitory effect of PQS during transcription. It is tempting to speculate that this suppression involves destabilizing non-B-DNA structures at PQS. Interestingly, genes up-regulated in BS tend to feature a region of increased PQS abundance 160-680 bp downstream of the TSS on the sense strand, overlapping a region 140-270 bp downstream of the TSS correlated with increased gene expression. It is difficult to account for this observation, and it suggests a possible BLM helicase-independent correlation between gene expression and PQS positioned 140-270 bp downstream of the TSS on the DNA sense strand.

**Figure 3.**
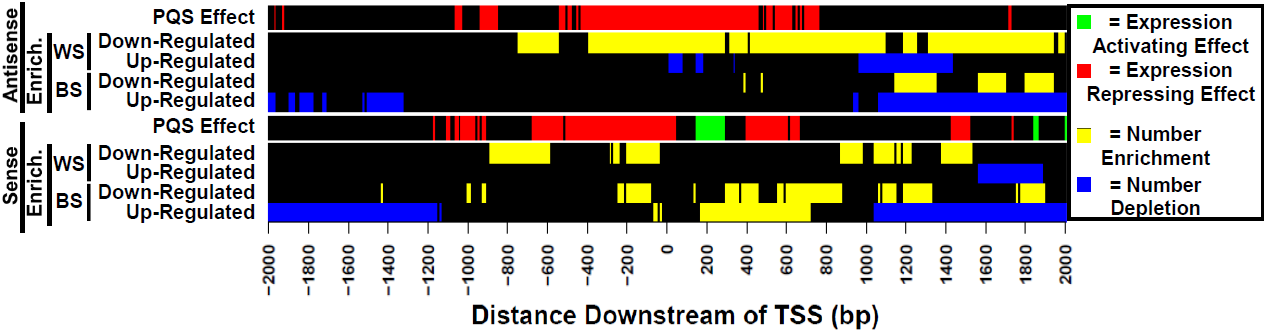
Heat map showing PQS excess or scarcity in genes differentially expressed in BS and WS (yellow = excess, blue = scarcity), as well as the correlation of PQS at these positions with transcription (green = high transcription, red = low transcription).

### PQS motifs in genes differentially expressed in BS and WS

To examine whether particular PQS motifs are correlated with transcription effects upon RecQ helicase loss, we analyzed PQS length, number of G-stacks, base composition, and loop lengths for genes differentially expressed in BS and WS, comparing these results to all other PQS. Loop lengths were calculated only for PQS containing four G-stacks, and lacking guanine at the first or last position of any loop sequence. PQS motifs in genes differentially-expressed in BS and WS were then compared to all other genes as a function of position upstream or downstream of the TSS (**Figure 4**). The results illustrate the interesting finding that PQS motifs in genes differentially expressed in BS and WS are statistically different from PQS in the remainder of the genome. Interestingly, these differences are consistent regardless of PQS position upstream or downstream of the TSS. FDR for this analysis was determined by statistical modeling using randomly-generated gene datasets, and is presented in **Supplemental Table S7A**. The position-independent similarity of PQS motifs in genes differentially expressed in BS and WS allowed an aggregate analysis of these motifs in comparison with all other PQS motifs near promoters of other genes (**Supplemental Table S7B**).

**Figure 4.**
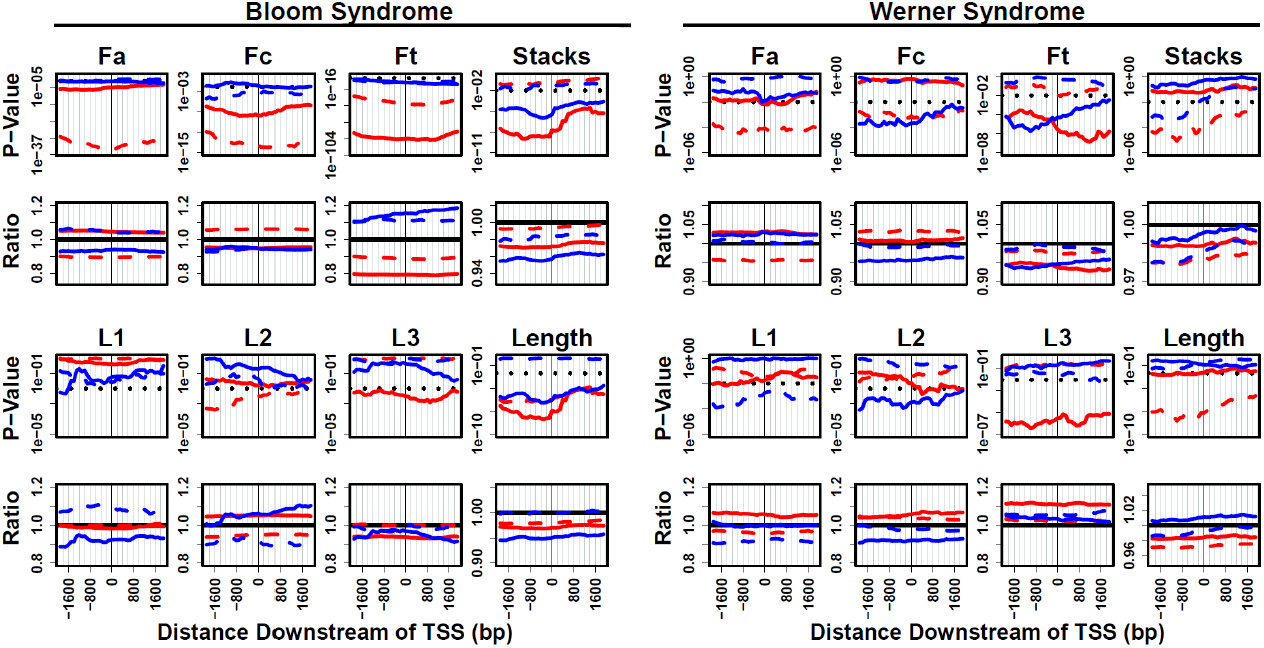
PQS motif characteristics (total sequence length, loop lengths, base fractions, and number of G-stacks) in genes differentially expressed in BS and WS. P-values and ratios for comparison to the genome-wide distribution are calculated using a 200 bp bin selection, repeated at a 50 bp interval, and in reference to corresponding PQS position on antisense (AS) strand or sense (S) strand for all genes. Red = genes up-regulated in BS and WS. Blue = genes down-regulated in BS and WS. Solid line = AS. Dashed line = S.

### Multidimensional PQS motif clustering

We further analyzed PQS motifs present in genes differentially expressed in BS and WS using a multidimensional self-organizing map (SOM) neural network classification. SOMs were historically designed to represent a high-dimensional space as a simple two-dimensional topological map (Kohonen 1982). We implemented SOM clustering of all PQS containing four G stacks. Using a 5X5 input matrix, SOM clustering was conducted using PQS total motif length, loop lengths, and base composition as the input dataset for training. PQS features for motif centroids of the 25 nodes resulting from the clustering protocol are presented in **Figure 5A**. The number of PQS counts per node, calculated on the basis of shortest multidimensional distance, is presented in **Figure 5B**. The multidimensional distance between nodes is represented in **Figure 5C**. To ascertain whether particular PQS motifs are associated with genes differentially expressed in BS and WS, we extracted these PQS motifs for these genes and calculated the average number of PQS per node, normalized to the total number of genes in the dataset. The same gene number-normalized calculation was repeated for the genome-wide dataset. A 95% confidence interval for normal statistical variability in cluster composition was also determined by statistical modeling conducted by repeating the same calculation on randomly-generated gene sets of the same size as the differential expression gene datasets. This comparison of PQS motif clusters from genes differentially expressed in BS and WS confirmed that certain PQS motifs are enriched in a statistically-meaningful fashion (**Figure 5D**). This provocative finding reveals a PQS motif bias in genes differentially expressed upon loss of RecQ helicases.

**Figure 5.**
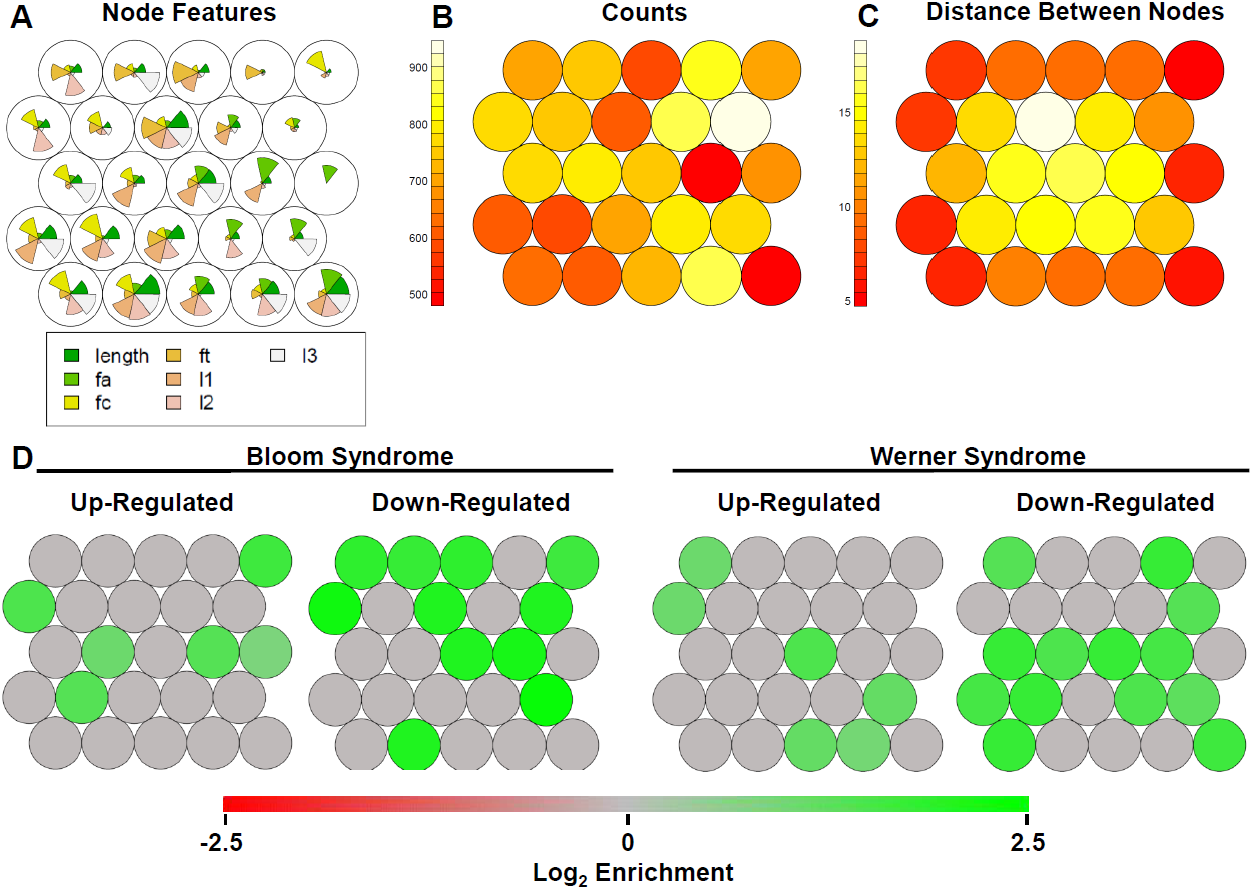
Multidimensional self-organizing map clustering of all human genome PQS within 2 kbp of TSS, based on total sequence length, loop lengths, number of G-stacks, and nucleotide base fractions. (A) PQS parameters for node centroids. (B) Counts of PQS binned by distance in multidimensional space to closest node. (C) Distance between nodes. (D) PQS node bias in genes differentially expressed in BS and WS. Bias represents number of unique PQS per node, normalized to dataset gene number, compared to the same calculation for all genes genome-wide. Green and red nodes represent positive (excess) or negative (scarcity) bias values that are outside of the 95% CI for randomly-generated gene datasets of the same size. Green = excess. Red = scarcity. Gray = no statistically-significant difference.

## DISCUSSION

We hypothesized that PQS patterns in genes differentially transcribed in BS and WS reveal transcriptional effects of PQS. In essence, BS and WS are natural RecQ helicase knockout experiments where transcriptional effects of persistent PQS non-B DNA structures are revealed. We further hypothesize that PQS in genes differentially-expressed in BS and WS are biased in their composition. We report analyses that support these hypotheses.

### Promoter PQS abundance correlates with transcriptional effect upon RecQ helicase loss

Regarding the first hypothesis, our analysis of PQS in promoters of genes differentially-expressed in BS and WS shows that patterns of PQS abundance in up-and down-regulated genes are opposite (**Figure 1**). Genes up-regulated upon RecQ helicase loss have a scarcity of promoter PQS and down-regulated genes have an abundance of PQS. This is reflected in the higher or lower numbers of PQS per TSS calculated for these gene sets compared to the genomic average (BS up-regulated: 1.61, BS down-regulated: 2.80, WS up-regulated: 1.58, WS down-regulated: 2.42, genome-wide average: 1.80; **Supplemental Table S3**). Interestingly, up-and down-regulated genes in BS and WS are more likely to have at least one TSS within 2 kbp of a PQS (BS: 84% of up-regulated genes and 90% of down-regulated genes, WS: 74% of up-regulated genes and 84% of down-regulated genes, genome-wide average: 55%; **Supplemental Table S1**). The implication of these results is that genes up-regulated in BS and WS have fewer PQS per TSS than the genomic average, but, surprisingly, are unlikely to have zero PQS motifs. Our results support a model in which promoter PQS generally repress transcription and RecQ helicases tend to moderate this repression.

### Promoter PQS abundance is altered at discrete positions in genes sensitive to RecQ helicase loss

Beyond PQS abundance, we find that PQS position is important. Thus, PQS abundance in genes differentially expressed in BS and WS is not randomly distributed across promoters, but discrete positions of abundance/scarcity are detected. The most striking example of this position-dependence is on the sense strand of genes up-regulated in BS (**Figure 1B**). Here PQS are scarce > 1 kbp upstream and downstream, but abundant 160-680 bp downstream of the TSS. This subtle and perplexing PQS position-dependence is missed when evaluating only aggregate PQS numbers per TSS.

Our analysis expands the current state of knowledge regarding PQS abundance in genes differentially expressed in BS and WS. Johnson et al. have shown that PQS are more abundant in non-coding regions of genes up-regulated in BS and WS (Johnson et al. 2010). This study, however, did not find statistically-significant correlation between PQS and genes down-regulated in BS and WS. Recent work improved upon this analysis by demonstrating increased PQS abundance in genes down-regulated in BS up to 250 bp upstream of the TSS on the antisense strand and at the 5’ end of the first intron on the sense strand (Nguyen et al. 2014). Genes up-regulated in BS were also found to have increased PQS abundance flanking the TSS (within 250 bp) on the antisense strand and at the 5’ end of the first intron on both sense and antisense strands. Interestingly, our analysis does not corroborate the findings of Nguyen et al. that PQS abundance is increased up to 250 bp upstream of the TSS on the antisense strand in genes down-regulated in BS, nor the finding that PQS abundance is increased flanking the TSS on the antisense strand for up-regulated genes. We suggest that the discrepancy is due to the rigorous statistical criteria in our analysis, and the higher resolution (smaller bin size) than was used by Nguyen et al. Thus, our data analysis examines PQS patterns further upstream and downstream of the TSS (a full 4 kbp window), and at a higher resolution (200 bp) than any previous study. We provide the first evidence that PQS abundance is significantly lower in the promoter-proximal region of genes up-regulated in BS and WS. The finding that PQS abundance is generally higher in genes down-regulated in BS and WS and lower in genes up-regulated genes in BS and WS, compared to the genomic average, provides an important insight. Again, these results support a model in which promoter PQS are generally repressive and RecQ helicases tend to modulate this repression.

### PQS position correlates with transcriptional effect genome-wide

We have also correlated PQS position with gene expression while controlling for epigenetic status to isolate intrinsic PQS transcriptional effects (**Figure 2**). For 33% and 35% of analyzed PQS positions in the antisense and sense strands, respectively, there was a statistically-significant correlation between PQS abundance and gene expression. Of these statistically-significant regions, 100% and 84% on antisense and sense strands, respectively were correlated with reduced expression. This suggests that transcriptional repression may be the dominant effect of promoter-proximal PQS, although certain PQS positions are correlated with increased gene expression (e.g. 140-270 bp downstream of TSS on the sense strand).

While there is general agreement with results of previous work, our genome-wide correlation of promoter PQS with gene expression provides a significant improvement over previous analyses. A prior analysis (Du et al. 2008) also correlated PQS in the sense strand up to 500 bp downstream of the TSS with higher gene expression. This prior analysis, which controls for attributes of gene family, function, and promoter similarity, did not find statistically significant correlation of PQS upstream of the TSS with gene expression. In contrast, the robust statistical analysis reported here, which controls for epigenetic status, shows that PQS upstream of the TSS on either DNA strand are correlated with lower gene expression. In addition to this greater sensitivity, our genome-wide correlation of promoter PQS and gene expression covers a larger genomic window (2 kbp upstream and downstream of each TSS) at a higher resolution (200 bp) than previously reported by Zhuo et al. (500 bp). For many of the genes differentially expressed in BS and WS, important PQS positions are > 500 bp distal from the TSS (86% for BS, 71% for WS). The study by Zhuo et al. was not able to provide insight into gene expression correlations for PQS in these positions.

Our simultaneous analysis of PQS abundance in genes differentially expressed in BS and WS, and our genome-wide correlation of promoter PQS with gene expression are unique. (**Figure 3**).

### PQS motifs are biased in genes sensitive to RecQ helicase loss

Regarding the second hypothesis of this work, we show that PQS motifs in genes differentially expressed in BS and WS are significantly biased in their composition (**Figure 4, Figure 5**). This indicates that specific sequence patterns may be important in PQS biological function. The reason for this is not clear, but it could reflect specificity in the PQS-helicase interaction. This is the first demonstration of PQS motif bias in a differentially-expressed gene set.

### PQS abundance and correlations with transcription suggest a hypothetical regulatory model

Interpreting these correlations between PQS and gene expression in the presence and absence of RecQ helicases is challenging. In the absence of biochemical evidence, caution is urged in attributing PQS effects to non-B DNA structures. Nonetheless, these data might be interpreted as evidence that non-B DNA G-quadruplex structures have position-dependent regulatory effects on gene expression, and these effects can be modified by RecQ helicases. In particular, the data support a model in which intrastrand G-quadruplexes generally inhibit transcription, with RecQ helicases tending to suppress this inhibition. Such a model would explain why loss of RecQ helicases results in decreased expression of genes with more promoter PQS, and greater expression of genes with fewer promoter PQS. Our data suggest a slightly more complex model in that increased sense strand PQS abundance 160-680 bp downstream of the TSS in genes up-regulated in BS (overlapping a region 140-270 bp downstream of the TSS correlated with increased gene expression) implies that PQS in these positions activate transcription in a BLM helicase-independent manner. A model to rationalize this observation could include sense strand PQS folding into intrastrand G-quadruplex structures at this position during transcription activation, enabling better access of the transcriptional machinery to the antisense strand. In this case BLM helicase interaction with the G-quadruplex might inhibit binding of other transcription-activating factors. BLM helicase loss would result in increased gene transcription in these special cases. Clearly, definitive structural interpretations of these data are beyond the scope of the present work.

## METHODS

### PQS identification

Putative intrastrand G-quadruplex-forming sequences matching the pattern G_3+_N_1-7_G_3+_N_1-7_G_3+_N_1-7_G_3+_ were identified within the GRCh37 build of the human genome using the Quadparser algorithm written in Python and described previously (Huppert and Balasubramanian 2005). All other data processing and statistical modeling was implemented in R version 3.0.2 run in a Windows 8.1 environment unless otherwise noted. From the output of the Quadparser program, reverse complement sequences for minus strand PQS were generated using the R package Biostrings available from the Bioconductor open source software project.

### PQS mapping near TSS

GENCODE version 7 gene annotations (produced using GRCh37) were downloaded from www.gencodegenes.org/releases/7.html. The list of annotations was filtered to remove all entries with a tag of “cds_start_NF” or “low_sequence_quality”, leaving a set of 156,253 GENCODE v7 transcripts. From this list of transcripts, 139,758 unique transcription start sites (TSS) were identified, representing 59,005 unique genes. PQS were then mapped to the 2 kbp upstream and downstream of identified TSS using PQS midpoint reference genomic coordinates. A total of 88,058 unique PQS were mapped to within 2 kbp of a TSS. Since many genes have multiple TSS located within a few kbp, some PQS were counted multiple times, with a total of 251,386 PQS mapping assignments to known TSS.

### BS and WS gene expression data

Differential gene expression data for Bloom Syndrome patient fibroblasts was obtained from the supplemental information section of reference (Nguyen et al. 2014). Genes in this dataset had been identified by Affymetrix GeneChip Human Exon 1.0 ST arrays using manufacturer-recommended protocols, with the criterion for differential expression set at ≥ 1.5 absolute fold expression change with an adjusted p-value < 0.05 and FDR < 0.1. Datasets for genes differentially expressed in Werner Syndrome patient fibroblasts were obtained from reference (Cheung et al. 2014) uploaded to GEO accession GSE48761. Genes in this dataset had been identified using Human Gene 1.0 ST Array (Affymetrix) using standard procedures. Datasets were downloaded from the GEO repository and processed in R using microarray analysis packages available from Bioconductor. Packages included hgu95av2cdf, hgu95av2.db, limma, marray, affy, affyQCReport, and affyPLM. WS datasets were normalized using the robust multiple-array average (RMA) algorithm, and genes differentially-expressed in WS patient samples were identified with the criterion of ≥ 1.5 absolute fold expression change compared to controls, with an adjusted p-value of < 0.05 and FDR < 0.05.

### Analysis of PQS in genes differentially expressed in BS and WS

Genes differentially-expressed in BS and WS were divided into up-and down-regulated gene sets. All TSS were identified in genes differentially-expressed in BS and WS, together with all PQS positioned within 2 kbp of these TSS. PQS were annotated for sense or antisense strand and positions assigned in 200 bp bins repeated at a 10 bp interval from −2 kbp to +2kbp relative to TSS. This approach was also applied to map all PQS in the human genome, and an additional 100 times in a statistical bootstrapping method using a randomly-generated collection of genes of the same size as the test dataset. Test datasets and randomly-generated datasets were compared to the genome-wide dataset using a p-value generated from the prop.test() function in R. P-values and the ratio of mean PQS per TSS for the comparison between the test datasets and genome-wide controls were plotted as a function of bin position relative to the TSS. The threshold for statistical significance was picked to be the p-value below which a data point in the randomly generated dataset has a 1% chance of false rejection of the null hypothesis. FRD were calculated as the ratio of predicted number of false positives data points (3.81 per 381 data points) to the number of data points in the test dataset that pass the threshold p-value.

### Epigenetic prediction of gene expression

The generation of predictive models for gene expression based on epigenetic data followed a method similar to that previously described (Dong et al. 2012). Epigenetic and gene expression data were obtained from the ENCODE project through the NCBI Gene Expression Omnibus online repository (accession GSE34448, GSE32970, and GSE29611). Cell lines used in the epigenetic modeling of gene expression include H1-hESC (embryonic stem cell), HeLa-S3 (cervical cancer), K562 (immortalized myelogenous leukemia), HUVEC (human umbilical vein endothelial cell), HepG2 (hepatocellular carcinoma), NHEK (normal human epidermal keratinocyte), and GM12878 (EBV-transformed lymphoblastoid cell). For each of these cell lines, genome-wide tracks for histone modifications H3K9ac, H3K4me3, H3K4me2, H3K27ac, H3K79me2, H3K36me3, H3K4me1, H3K27me3, H4K20me1, histone variant H2A.Z, and chromatin accessibility (quantified by digital DNAse I hypersensitivity) were obtained, along with gene expression datasets quantified by cap analysis of gene expression (CAGE) technology.

GENCODE version 7 transcript annotations were used to map individual CAGE-quantified transcript levels to known TSS. Total transcriptional activity for each TSS was calculated as the sum of the expression values for all transcripts that originate from that TSS. Since many genes contain multiple TSS, only the TSS with the highest aggregate expression level for each of the 59,005 unique genes in the genome was retained for analysis.

The GC fraction for the 5 kbp upstream of each of the identified strongest TSS was calculated using tools in the R package Biostrings produced by the open source Bioconductor software project. Computation for this part of the analysis was run on the Mayo Research Computing Facility (RCF) shared-resource, Beowulf-style Linux cluster using R version 3.0.2, with scripts written to accommodate batch mode execution managed by the Open Grid Engine open-source batch-queuing system.

ENCODE data for DNAse I hypersensitivity, histone modification, and histone variant tracks were obtained from the GEO repository as described above. Track signal files were downloaded in .bigwig format and processed using utilities in the R package rtracklayer from Bioconductor. Track signals within 1 kbp upstream and downstream of each TSS were extracted and the signal within this region was split and averaged into twenty 100-bp bins, spanning from 1 kbp upstream to 1 kbp downstream of each TSS. For each of the 20 bins constructed for each genomic track, a correlation coefficient was calculated between the log_2_-transformed bin signal (with a small pseudocount added to avoid the log_2_(0) issue) and log_2_-transformed gene expression signal quantified via CAGE, excluding TSS with a expression value of 0. For each genomic feature, the bin that had the highest correlation with expression was selected as the “best bin” for analysis. The optimal pseudocount value for each genomic track was determined by repeating the correlation analysis described above, but with pseudocounts ranging from 0.25-5% of the maximal binned signal for that track feature. For each genomic track, the best pseudocount and “bestbin” combination was selected by identifying the bin and pseudocount combination that resulted in the highest absolute correlation with gene expression.

“Bestbin” and pseudocount combinations for each epigenetic track, along with the GC% 5 kbp upstream of each TSS were then used to construct predictive models of gene expression. TSS with missing values for any of the genomic tracks were excluded, leaving between 4,011 and 9,614 TSS, depending on the cell line. The subset of TSS containing no PQS within 2 kbp of the TSS was then divided into equal training and test datasets using a random sampling approach. The subset of TSS containing at least one PQS was included in the test dataset. All data channels in training and test datasets were normalized to be mean-centered at 0 and to have a standard deviation of 1. A Bayesian linear regression model for predicting gene expression from epigenetic parameters and GC fraction was then generated using the training dataset and the R package tgp. This model was then applied to the test dataset (not used in the training of the model) to generate gene expression predictions.

### Modeling PQS correlation with transcription

For each TSS in the epigenetic modeling of gene expression test dataset, a prediction error was calculated as the CAGE-determined TSS-specific expression value minus the Bayesian linear regression model prediction from the epigenetic data. The portion of the test dataset TSS containing at least one PQS was extracted and the position and strand of the nearest PQS to each TSS was determined. Using an iterative approach, TSS-specific prediction error data for sense and antisense strands were aggregated based on the location of the nearest PQS, using a 200 bp bin size, and running 2 kbp upstream to 2 kbp downstream at an iteration interval of 10 bp. For each bin, the distribution of prediction errors was compared to the prediction errors of the TSS in the test dataset containing no PQS using the oneway.test() function in R for one-way ANOVA. As a statistical control to assess the likelihood of obtaining a low p-value in the comparison by random chance, a statistical bootstrapping approach was employed in which the prediction errors in the PQS dataset were replaced with random prediction errors from the test dataset, and the p-values were calculated as above. This comparison was repeated 100 times and the threshold p-value for determining statistical significance of the test dataset comparison was picked to be the p-value below which a data point in the control dataset has a 1% chance of false rejection of the null hypothesis. P-values in the test dataset below this threshold value were selected as statistically significant. FDR were calculated as the ratio of significant p-value data points in the test dataset compared to the average number of false positives in the control datasets (3.81 false discoveries per 381 data points).

### PQS motif analysis

The number of G stacks containing at least 3 contiguous guanine nucleotides in each PQS within 2 kbp of a TSS was calculated for all human genes using a pattern-matching algorithm. The nucleotide fractions of adenine, thymine, and cytosine in each PQS were calculated using the package Biostrings. For the subset of PQS containing 4 G stacks, with each G stack containing 3 bases, lengths of loop sequences between G stacks were also calculated. In order to avoid ambiguity in this analysis, it was required that the first and last base in a loop sequence not be guanine. Computation was performed on the Mayo Research Computing Facility (RCF) shared-resource, Beowulf-style Linux cluster using R version 3.0.2, with scripts written to accommodate batch mode execution managed by the Open Grid Engine open-source batch-queuing system.

### PQS motif analysis in genes differentially expressed in BS and WS

PQS were binned based on position, using a 200 bp bin width calculated at a 50 bp interval, spanning from 2 kbp upstream to 2 kbp downstream of the TSS. Subsets of PQS data for genes up-and down-regulated in BS and WS were extracted, and PQS features (total length, loop lengths, number of G-stacks, fractions adenine, cytosine, and thymine) were compared individually between genes differentially expressed in BS and WS and the remainder of the genome, bin by bin, using one-way ANOVA. Comparisons were done independently for PQS in the sense and antisense DNA strands. This bin-based comparison was repeated 100 times in a statistical bootstrapping method using a randomly-generated collection of genes of the same size as the test dataset. The threshold for statistical significance was picked to be the p-value below which a data point in the randomly generated dataset has a 1% chance of false rejection of the null hypothesis. False discovery rates were calculated as the ratio of predicted number of false positives data points to the number of data points in the test dataset that pass the threshold p-value.

### Self-organizing map multidimensional classification of PQS

From the entire human genome, the subset of PQS with 4 G-stacks and 3 loops within 2 kbp of a TSS was selected, excluding PQS in which the first or last base in a loop was guanine. This selection criterion reduced the number of surveyed PQS from 88,058 to 17,795. Each PQS in this selected dataset was classified for total sequence length, loop lengths, and fractions of adenine, cytosine, and thymine. Self-organizing map artificial neural network analysis was implemented in R using the package kohonen. Maps consisting of 25 nodes in a hexagonal grid were trained using the PQS dataset, 100 iterative presentations of the data to the model, and a learning rate with linear decline from 0.05 to 0.01. The average number of PQS per gene per node was calculated and compared to the same calculation repeated for the PQS and gene subset for genes up-and down-regulated in BS and WS. Log_2_ enrichment ratios were calculated for each node in the BS and WS datasets. As a means to ascertain whether the enrichment of PQS in certain nodes could arise by chance, a statistical bootstrapping method was employed in which enrichment ratios of nodes on the PQS map were calculated for 100 randomly-generated gene sets of the same size as the BS and WS datasets. Mean and 2σ values were calculated node by node for these random distributions. Statistical significance for enrichment ratios of nodes in the BS and WS datasets was assigned on the basis of lying outside of the 95% CI for the random distributions.

## DATA ACCESS

Supplemental datasets and code in support of this manuscript may be accessed at ftp://www.jsmes.net/PQS_Genomics.

## ACKNOWLEDGMENTS

We wish to thank Dr. Justin Peters for his technical advice relating to the use of the Mayo Research Computing Facility Linux cluster.

